# Olanzapine leads to nonalcoholic fatty liver disease through the apolipoprotein A5 pathway

**DOI:** 10.1101/2021.03.02.433415

**Authors:** Rong Li, Wenqiang Zhu, Piaopiao Huang, Yang Yang, Fei Luo, Wen Dai, Li Shen, Wenjing Pei, Xiansheng Huang

**Author notes:** Correspondence: Xiansheng Huang, Department of Cardiovascular Medicine, The Second Xiangya Hospital, Central South University, #139 Middle Renmin Road, Changsha, Hunan 410011, China.

## Abstract

The antipsychotic drug olanzapine was reported to induce nonalcoholic fatty liver disease (NAFLD), whereas the underlying mechanism remains incompletely understood. This study investigated whether apolipoprotein A5 (apoA5) and sortilin, two interactive factors involved in NAFLD pathogenesis, are implicated in olanzapine-induced NAFLD. In the present study, at week 8, olanzapine treatment successfully induced hepatic steatosis in male C57 BL/6J mice, even when fed a normal chow diet, which was independent of animal body weight gain. Likewise, olanzapine effectively mediated hepatocyte steatosis in human HepG2 cells *in vitro*, characterized by substantially elevated intracellular lipid droplets. Increased plasma triglyceride concentration and decreased plasma apoA5 levels were observed in mice treated for 8 weeks with olanzapine. Surprisingly, olanzapine markedly enhanced hepatic apoA5 protein levels in mice, without a significant effect on rodent hepatic *ApoA5* mRNA expression. Our *in vitro* study showed that olanzapine reduced apoA5 protein levels in the medium and enhanced apoA5 protein expression in hepatocytes, whereas this drug exerted no effect on hepatocyte *APOA5* mRNA expression. By transfecting *APOA5* siRNA into HepG2 cells, it was demonstrated that *APOA5* knockdown effectively reversed olanzapine-induced hepatocyte steatosis. In addition, olanzapine drastically increased sortilin mRNA and protein levels *in vivo* and *in vitro*. Interestingly, *SORT1* knockdown reduced intracellular apoA5 protein expression and increased medium apoA5 protein levels *in vitro*, without affecting intracellular *APOA5* mRNA expression. Furthermore, *SORT1* knockdown greatly ameliorated hepatocyte steatosis *in vitro*. This study provides the first evidence that sortilin inhibits the hepatic apoA5 secretion that is attributable to olanzapine-induced NAFLD, which provides new insight into effective strategies against NAFLD for patients with schizophrenia administered olanzapine.

## 1. Introduction

Nonalcoholic fatty liver disease (NAFLD), a pathological state associated with the excessive accumulation of triglycerides in the liver without significant alcohol consumption [1], is the leading cause of chronic liver disease, with a global prevalence rate as high as 25.34% [2]. This encompasses a broad spectrum of conditions from simple steatosis to non-alcoholic steatohepatitis that can progress to cirrhosis and hepatocellular carcinoma [3, 4]. Although the pathogenesis of NAFLD is mostly attributed to overnutrition-associated obesity, this disease might partly be caused by side effects of pharmacological therapies. Specifically, long-term treatment with olanzapine, a representative second-generation antipsychotic drug widely used for patients with schizophrenia, is strongly associated with obesity and metabolic syndrome [5–8], both of which are predisposing factors for NAFLD. Several studies have reported that olanzapine promotes the development of NAFLD in rodents [9, 10]. However, the mechanism underlying olanzapine-induced NAFLD has not been fully elucidated.

Indeed, NAFLD and obesity are strongly associated with disorders of triglyceride homeostasis, both of which develop when the triglyceride production rate exceeds the catabolic rate in the liver and adipose tissue, respectively. Triglycerides represent the major storage and transport forms of fatty acids, and the liver is the central organ for fatty acid metabolism. Physiologically, the liver stores only small amounts of fatty acids, which are localized in lipid droplets within hepatocytes. In the setting of overnutrition, hepatic fatty acid metabolism is disturbed, which results in the excessive hepatic accumulation of triglycerides and eventually the development of NAFLD.

Apolipoprotein A5 (apoA5), a liver-specific synthesized and secreted protein [11, 12], is an essential regulator of triglyceride metabolism *in vivo* [11]. It has been well documented that apoA5 plays a crucial role in reducing plasma triglyceride levels [13, 14]. Interestingly, *APOA5* genetic polymorphisms are associated with olanzapine-mediated dyslipidemia [20, 21], suggesting that apoA5 might participate in this process. Of note, apoA5 is also involved in the pathogenesis of NAFLD. Increased hepatic *APOA5* mRNA expression was found in obese-associated adult [22] and pediatric [23] NAFLD livers, whereas reduced hepatic *APOA5* mRNA expression was detected after weight loss and NAFLD improvement [21]. Increased *ApoA5* mRNA expression has also been observed in rodent NAFLD livers [22]. Compared to those in wild-type and *ApoA5*-knockout mice, *ApoA5* transgenic mice have higher levels of hepatic triglycerides, which results in hepatic steatosis in rodents [24]. Furthermore, apoA5-induced intracellular lipid accumulation leads to a substantial increase in hepatic lipid droplets [24–27], a critical pathological hallmark of NAFLD. Therefore, these data support a role for apoA5 as a potential target for NAFLD. To date, however, it remains unknown whether apoA5 mediates olanzapine-induced NAFLD.

Sortilin, encoded by the *SORT1* gene on chromosome 1p13.3, is a multi-ligand transmembrane protein that traffics newly synthesized molecules from the trans-Golgi apparatus along secretory pathways to endosomes, lysosomes, or the cell surface. Previous studies have established an essential role for hepatic sortilin in lipoprotein transport and secretion in the liver [28, 29]. Recently, it has been shown that increased hepatic sortilin expression markedly enhances susceptibility to NAFLD [30–32]. Specifically, sortilin localizes to the plasma membrane in clathrin-coated pits where it can function as an internalization receptor for apoA5, after which apoA5 rapidly internalizes and co-localizes with sortilin in early endosomes, followed by sortilin trafficking through endosomes to the trans-Golgi apparatus [33]. These data strongly suggest that sortilin might be implicated in apoA5-mediated hepatic lipid accumulation; this and the role of sortilin in olanzapine-induced NAFLD was investigated in this study.

## 2. Methods

### 2.1. Animal studies

#### 2.1.1. Animals

All experimental procedures were carried out according to the recommendations of the National Institutes of Health Guide for the Care and Use of Laboratory Animals (NIH Publications No.8023, revised 1978) and were approved by the Experimental Animal Ethics Committee of the Second Xiangya Hospital, Central South University. Male C57 BL/6J mice (6-weeks-old, n = 36, Hunan Stryker Jingda Animal Co., Ltd) were used in the experiments. After 1 week of environmental acclimatization, animals were maintained in an air-conditioned room at 22–24°C with a 12-h light/dark cycle with *ad libitum* access to water. Mice were fed with a normal chow diet (10% fat, 70% carbohydrates, and 20% protein; D12450B, Research Diets) throughout the study.

#### 2.1.2. Drug treatment

Animal fatty liver was induced as previously described [34–36] with slight modifications. Briefly, mice were gavaged with olanzapine at a dose of 3 or 6 mg/kg body weight per day. Olanzapine (LY170053, MCE, USA) used for mice was prepared freshly prior to use and dissolved in 10% dimethyl sulfoxide (DMSO).

The mouse administration plan was established as follows. First, C57 BL/6J mice (n = 36) were randomly assigned into three groups (n = 12/each group). One group was subjected to daily gavage of 10% DMSO as the control group, whereas the other two groups received daily gavage of olanzapine (3 or 6 mg/kg body weight). Second, each of the three groups was randomized into two subgroups, specifically the 4- and 8-week subgroups (n = 6/each subgroup).

The body weights of animals were evaluated weekly throughout the study. Animals were fasted for 6 h and then anesthetized with pentobarbital at week 4 or 8, respectively. Blood samples were collected to detect plasma lipid levels (triglycerides, high-density lipoprotein cholesterol [HDL-C], low-density lipoprotein cholesterol [LDL-C], and total cholesterol) and to evaluate liver function according to the serum levels of alanine transaminase (ALT) and aspartate aminotransferase (AST). The liver samples were immediately dissected and frozen in liquid nitrogen and stored in a −80°C freezer until further analysis.

### 2.2 Cell studies

#### 2.2.1. Cell culture

The human hepatoma cell line (HepG2) was obtained from the American Type Culture Collection and cultured in standard medium comprising DMEM (Dulbecco’s modified Eagle’s medium), 10% FBS, and 1 % penicillin-streptomycin in an atmosphere containing 5% CO_2_ at 37°C. All cell lines in our laboratory were passaged 2–3 times per week and no more than 30 times after resuscitation.

#### 2.2.2. Drug treatment

Olanzapine (LY170053, MCE, USA) was dissolved in pure DMSO (D2650, Sigma, USA) and then added to the medium for treatment for 24 h at the final concentrations of 25, 50, and 100 μmol/l. DMSO was adjusted to 0.1% in the culture medium. The group treated with 0.1% DMSO served as a blank control.

### 2.3. Transient transfection

*APOA5* siRNA, *SORT1* siRNA, and their nonspecific siRNAs were constructed by Bio-Gene (stB0005146, Bio-Gene, China). Transfection of siRNAs was performed with Lipofectamine 3000 (L3000015, Thermo Fisher Scientific, USA) or Lipofectamine RNAiMAX (Thermo Fisher Scientific). Briefly, 6 × 10^5^ cells were seeded in six-well plates with 1.5 ml culture medium without antibiotics. Simultaneously, siRNAs were mixed with Lipofectamine 3000 and incubated at room temperature for 15 min. The complexes were then transfected into HepG2 cells in an incubator at 37°C with 5% CO_2_ for 48 h.

### 2.4. Oil red O staining

Frozen liver sections were subjected to Oil Red O staining (ORO staining) to visualize lipid droplets. The tissues were cut into 6-µm slices and then stained for lipids according to the standard ORO staining protocol (ORO, Merck, Germany). The samples were rinsed with 60% isopropanol and stained with filtered ORO solution (0.5% in isopropanol followed by 60% dilution in distilled water) for 30 min. After washing twice with 60% isopropanol and distilled water, the slides were counterstained with hematoxylin. All images were acquired using a microscope. The area of positive staining for ORO was calculated as a percentage of the total section area, and the average lipid droplet size was calculated using the morphometry software Image J (version 1.8.0) [9].

ORO staining was also used to assess intracellular lipid droplets. Briefly, HepG2 cells were washed twice with PBS (phosphate-buffered saline), fixed with 4% formaldehyde solution for 0.5 h at 37°C, and then washed three times with PBS. ORO staining solution was prepared as follows: the stock solution was made of 0.5 g of ORO powder (O0625, Sigma, USA) dissolved in 100 ml of 100% isopropanol, and then, it was diluted with water (6:4) and filtered. Staining was conducted for 30 min at 37°C. Subsequently, the plates were destained in 60% isopropanol, and ORO staining was visualized using phase-contrast microscopy. For quantitative analyses of the ORO content, 100% isopropanol was added to each well to extract the ORO combined with lipid droplets. The absorbance was measured at 520 nm and then analyzed using GraphPad Prism software [37].

### 2.5. Hematoxylin and eosin staining

Hematoxylin and eosin (H&E) staining was performed to observe the distribution of accumulated lipids in liver tissue. The samples were fixed in 10% neutral formalin at 4°C overnight, dehydrated in ethanol, and then embedded in paraffin. Tissue sections (6-μm) were cut and mounted on glass slides for H&E staining per standard procedures [38].

### 2.6. Western blotting analysis

Total protein samples were isolated from liver tissue or cells using RIPA lysis buffer (P0013B/D, Beyotime Biotechnology, China) containing 1% 0.5 mmol/l phenylmethanesulfonyl fluoride. After incubation and centrifugation, the supernatant was saved as a protein extract. A vacuum freezing dryer (SCIENTZ-10N, Ningbo Scientz Biotechnology Co., Ltd.) was used for the concentration and purification of proteins in cell culture medium.

The protein levels were quantified using a micro BCA protein assay kit (CW0014S, CWBIO, China). The protein extracts were diluted with loading buffer and PBS and then degenerated. Equivalent amounts of protein samples were subjected to SDS/PAGE electrophoresis, and proteins were transferred onto a PVDF membrane. After the non-specific binding sites were blocked with 5% skim milk, the membranes were incubated in a blocking buffer containing the appropriate diluent of primary antibodies against apoA5 (#3335, Cell Signaling Technology, USA; sc-373950, Santa Cruz, USA), tubulin (66031-1-lg, Proteintech, USA), and sortilin (ab16640, Abcam, England) at 4°C overnight. After that, the membranes were washed and incubated with the corresponding HRP-conjugated secondary anti-mouse and anti-rabbit antibodies (SA00001-2, Proteintech, USA; SA00001-1, Proteintech, USA) for 2 h and then diluted 1:5000 in 5% milk/-PBST. Finally, the protein bands were visualized using Pierce^™^ ECL western blotting Substrate (32209, Thermo Scientific, USA) and quantified using a Molecular Imager ChemiDoc^™^ XRS+ (Bio-Rad). Densitometry analysis was performed using Image J v1.8.0.

### 2.7. Quantitative real-time PCR (qPCR) analysis

For qPCR analysis, total mRNA was isolated from liver tissues or cells using an RNA extraction kit (K0731, Thermo Scientific, USA). The cDNA was synthesized from 2 μg of purified RNA using the Revert Aid First Strand cDNA Synthesis Kit (K1622, Thermo Scientific, USA). SYBR Green (K1622, Thermo Scientific, USA) was used to quantify PCR amplification products. The mRNA expression levels of the target genes were normalized relative to those of the endogenous gene *GAPDH*. The primer pairs (synthesized by Tsingke Biological Technology) used in this study are listed in Table 1.

**Table 1.**
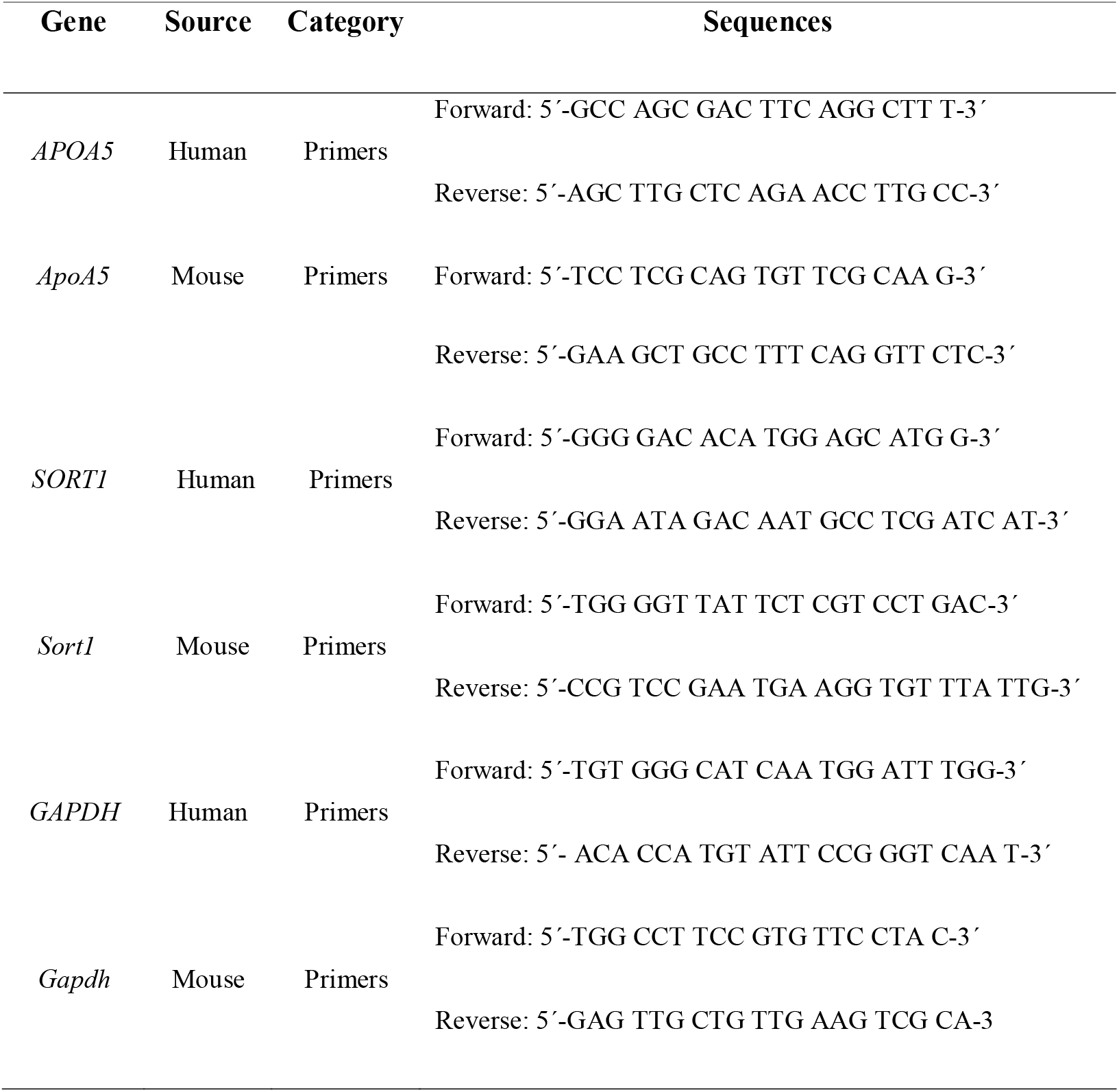
**Oligonucleotide sequences of primers targeting human and mouse genes**

### 2.8. Statistical analysis

All data were analyzed using the appropriate statistical analysis methods, as specified in the figure legends, with GraphPad Prism software (version 8.0). Statistical differences in measures of the different groups were analyzed by Student’s two-tailed t-tests (two groups) and one-way analysis of variance followed by Tukey’s multiple-comparison post hoc test (more than two groups). Statistical significance was set at *p* < 0.05. All results are presented as the mean ± SEM. No data were excluded when conducting the final statistical analysis. Randomization and blinding were performed when grouping mice and collecting data.

## 3. RESULTS

### 3.1. Effect of olanzapine on body weight and liver weight in animals

We monitored the olanzapine-induced changes in body weight in mice from baseline to 14 weeks (Fig 1A). Interestingly, there was no significant difference in body weight among the three groups during the study, suggesting that olanzapine treatment (3 and 6 mg/kg) exerted no pronounced effect on rodent body weight within a certain short period (≤ 8 weeks). However, olanzapine dose- and time-dependently increased liver weights in mice (Fig 1B). Similarly, olanzapine remarkably increased the liver weight/body weight ratio in mice in a dose- and time-dependent manner (Fig 1C). Taken together, these data show that olanzapine-induced liver weight gain in mice is independent of body weight gain.

**Figure 1.**
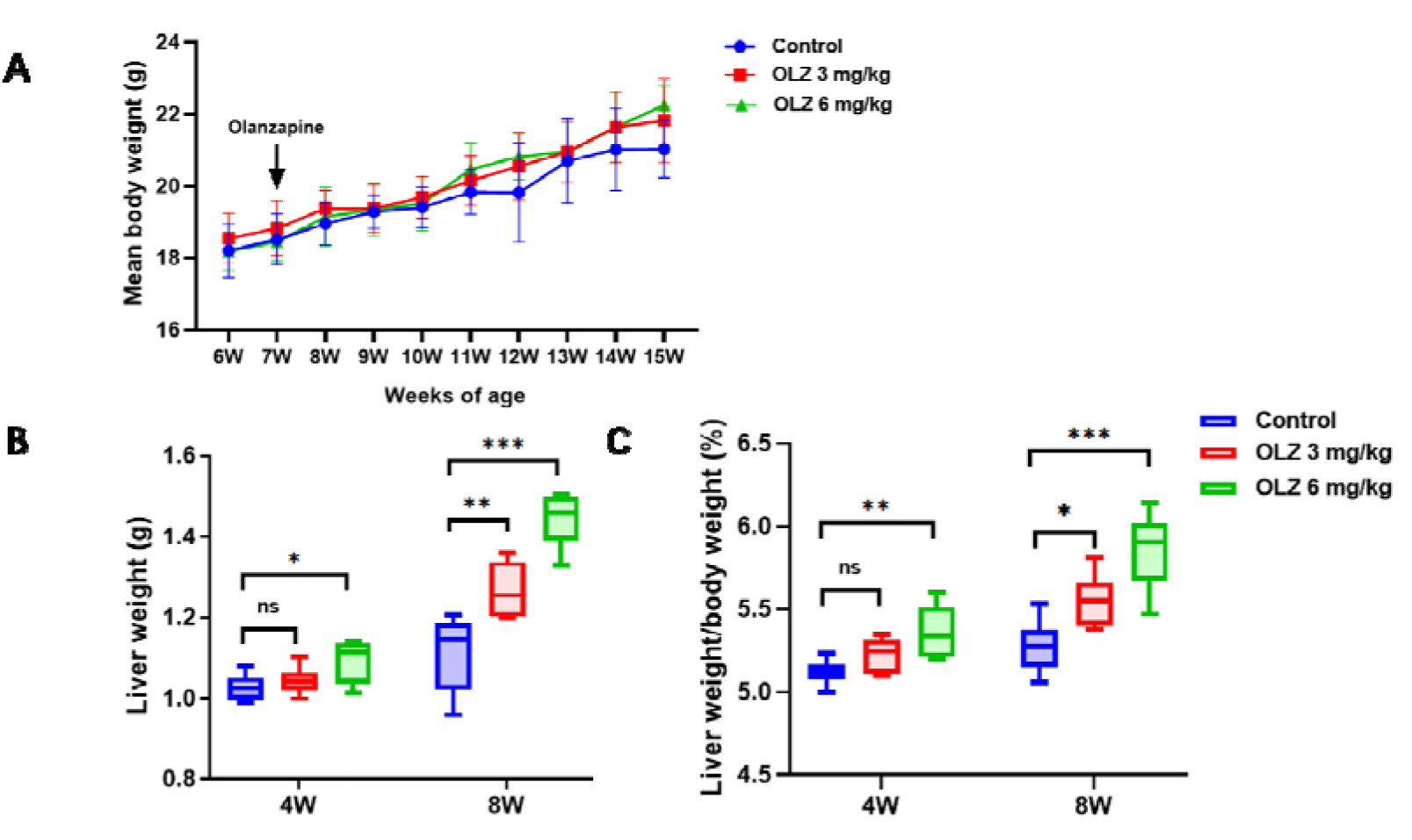
Effect of olanzapine on body weight and liver weight in mice (n = 6/each group). (**A**) Body weight. (**B**) Liver weight. (**C**) Liver weight/body weight (LW/BW) ratio. OLZ, olanzapine. Results are shown as the mean ± SEM. *\*p* < 0.05, *\*\*p* < 0.01, *\*\*\*p* < 0.001, versus the control group. One-way ANOVA plus Tukey’s test was performed for the data analysis.

### 3.2. Olanzapine increases plasma triglycerides and hepatic steatosis in mice and hepatocyte triglycerides in vitro

Plasma lipid levels in mice were detected at weeks 4 and 8 (Table 2). There was no significant difference in the plasma levels of all lipids at week 4, indicating that 4-week olanzapine treatment cannot lead to dyslipidemia in these animals. In contrast, higher plasma triglyceride levels were observed in the two olanzapine groups than in the control group at week 8, whereas higher plasma triglyceride levels were found in the 6 mg/kg olanzapine group (1.05 ± 0.18 mM) than in the 3 mg/kg olanzapine group (0.69 ± 0.16 mM, *p* < 0.05) at week 8, suggesting that olanzapine dose-dependently increased plasma triglyceride levels in mice at week 8. However, no significant difference was observed in the other three lipids (HDL-C, LDL-C, and total cholesterol) among the three groups at week 8. Thus, these data show that 8-week olanzapine treatment has a more detrimental effect on the metabolism of circulating triglycerides than other lipids in these mice.

**Table 2.** **Plasma lipid levels and liver enzymes of mice**

| Group | NEFA<br>(mmol/l) | Triglycerid<br>es (mmol/l) | Total<br>cholesterol<br>(mmol/l) | HDL-C<br>(mmol/l) | LDL-C<br>(mmol/l) | ALT<br>(U/l) | AST<br>(U/l) | AST/ALT |
| --- | --- | --- | --- | --- | --- | --- | --- | --- |
| **4W** |  |  |  |  |  |  |  |  |
| Control | 0.73 ± 0.18 | 0.38 ± 0.09 | 1.69 ± 0.18 | 1.41 ± 0.13 | 0.30 ± 0.05 | 26.43 ± 3.66 | 90.48 ± 8.88 | 3.47 ± 0.47 |
| 3 mg/kg<br>olanzapine | 0.82 ± 0.19 | 0.32 ± 0.04 | 1.71 ± 0.28 | 1.42 ± 0.21 | 0.30 ± 0.07 | 25.71 ± 5.57 | 105.60 ± 11.54 | 4.23 ± 0.77 |
| 6 mg/kg<br>olanzapine | 0.85 ± 0.27 | 0.31 ± 0.09 | 2.00 ± 0.16 | 1.71 ± 0.16 | 0.33 ± 0.05 | 28.43 ± 3.43 | 131.22 ± 47.73 | 4.59 ± 1.46 |
| **8W** |  |  |  |  |  |  |  |  |
| Control | 0.67 ± 0.16 | 0.39 ± 0.13 | 1.60 ± 0.20 | 1.45 ± 0.16 | 0.36 ± 0.05 | 27.30 ± 3.11 | 89.77 ± 6.50 | 3.35 ± 0.54 |
| 3 mg/kg<br>olanzapine | 0.79 ± 0.11 | 0.69 ± 0.16* | 1.61 ± 0.21 | 1.44 ± 0.19 | 0.28 ± 0.06 | 61.83 ± 14.87* | 114.89 ± 12.46* | 1.93 ± 0.37** |
| 6 mg/kg<br>olanzapine | 0.88 ± 0.19 | 1.05 ± 0.18** | 2.17 ± 0.27** | 1.67 ± 0.13 | 0.33 ± 0.03 | 106.05 ± 27.55** | 164.45 ± 23.38** | 1.63 ± 0.38** |

\* $p < 0.05$ , \*\* $p < 0.01$ vs. control
NEFA: nonesterified fatty acids; HDL-C: high-density lipoprotein cholesterol; LDL-C: low-density lipoprotein cholesterol;

Liver function markers (ALT and AST) in mice were next determined (Table 2). Similar to the aforementioned results of lipids, no significant difference was detected in serum aminotransferase levels among the three groups at week 4, which suggests that 4-week olanzapine treatment does not impair liver function in these mice. At week 8, however, we observed a markedly detrimental effect of olanzapine on rodent liver function in a dose-dependent manner. To investigate the effect of olanzapine on hepatic steatosis, we further detected hepatic triglyceride content by H&E (Fig 2A) and ORO staining (Fig 2B). Our data showed that olanzapine administration resulted in marked rodent hepatic steatosis in a dose-dependent manner. Likewise, our *in vitro* study using ORO staining demonstrated that olanzapine dose-dependently increased intracellular triglyceride contents in hepatocytes (Fig 2C). Together, these findings reveal that olanzapine treatment promotes the development of NAFLD.

**Figure 2.**
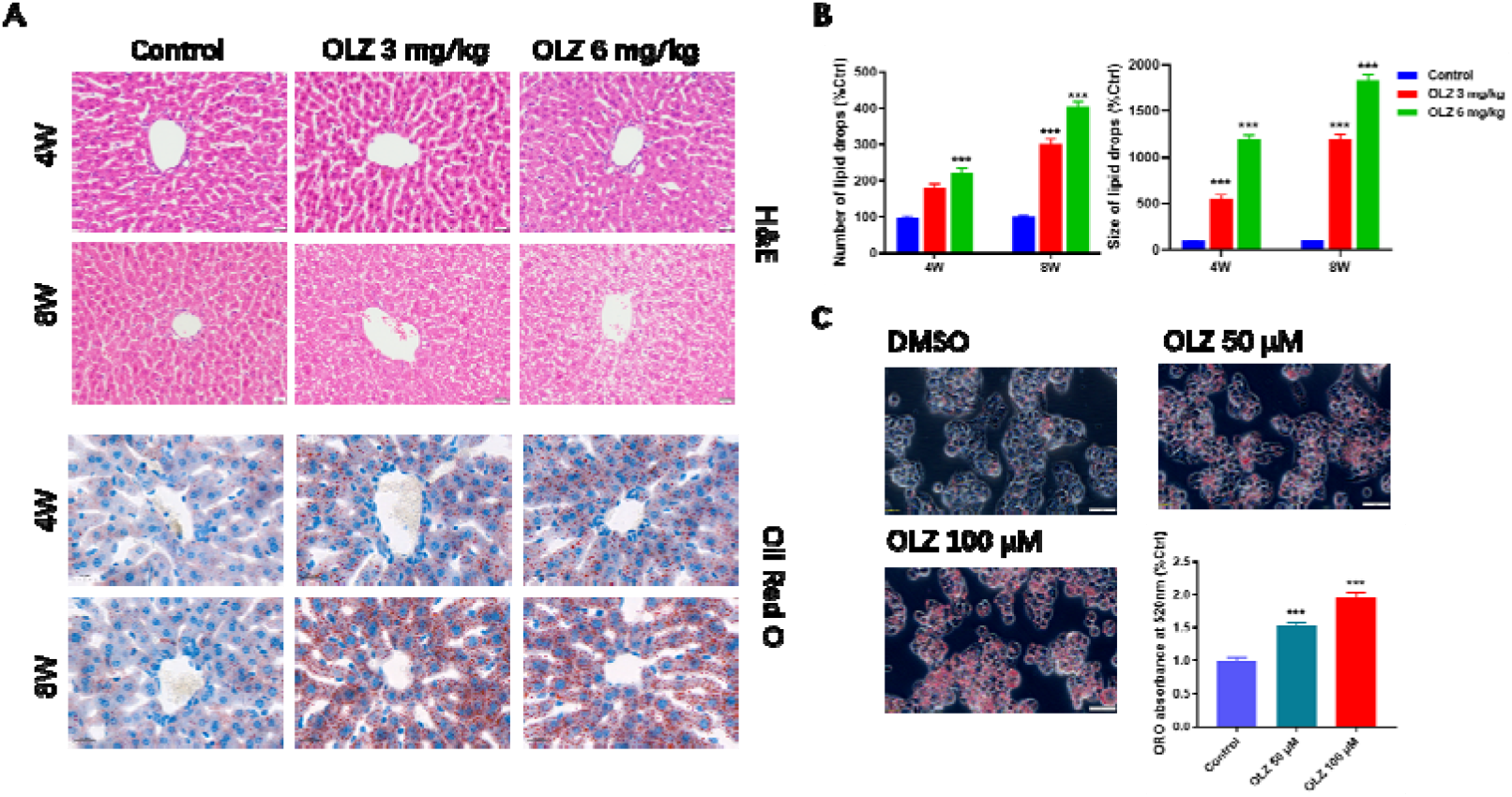
Effect of olanzapine on plasma lipid levels and hepatic/hepatocyte steatosis. (**A**) H&E and Oil Red O (ORO) staining of mouse liver tissue. (**B**) Number and size of hepatic lipid drops in mice by ORO staining. (**C**) ORO staining and absorbance in HepG2 cells. OLZ, olanzapine. Results are shown as the mean ± SEM. *** *p* < 0.001 versus control group. One-way ANOVA plus Tukey’s test was performed for the data analysis.

### 3.3. Olanzapine inhibits hepatic apoA5 secretion and exacerbates hepatic steatosis

To investigate the role of apoA5 in olanzapine-induced NAFLD, we detected plasma and hepatic apoA5 levels in mice. Our ELISA tests showed that olanzapine treatment markedly reduced plasma apoA5 levels (Fig 3A). Given that apoA5 is synthesized in a liver-specific manner and subsequently secreted into circulation [11,12], we further determined hepatic apoA5 protein and mRNA levels by western blotting and qPCR analyses, respectively. Surprisingly, olanzapine dose- and time-dependently increased hepatic apoA5 protein levels in mice (Fig 3B), whereas no significant difference was observed in hepatic *ApoA5* mRNA expression among the three groups (Fig 3B). Consistent findings were obt ained *in vitro*. Olanzapine drastically reduced apoA5 protein levels in the medium (Fig 3C), whereas it substantially enhanced apoA5 protein levels in hepatocytes (Fig 3D). However, olanzapine treatment did not influence *APOA5* mRNA expression in hepatocytes (Fig 3D).

**Figure 3.**
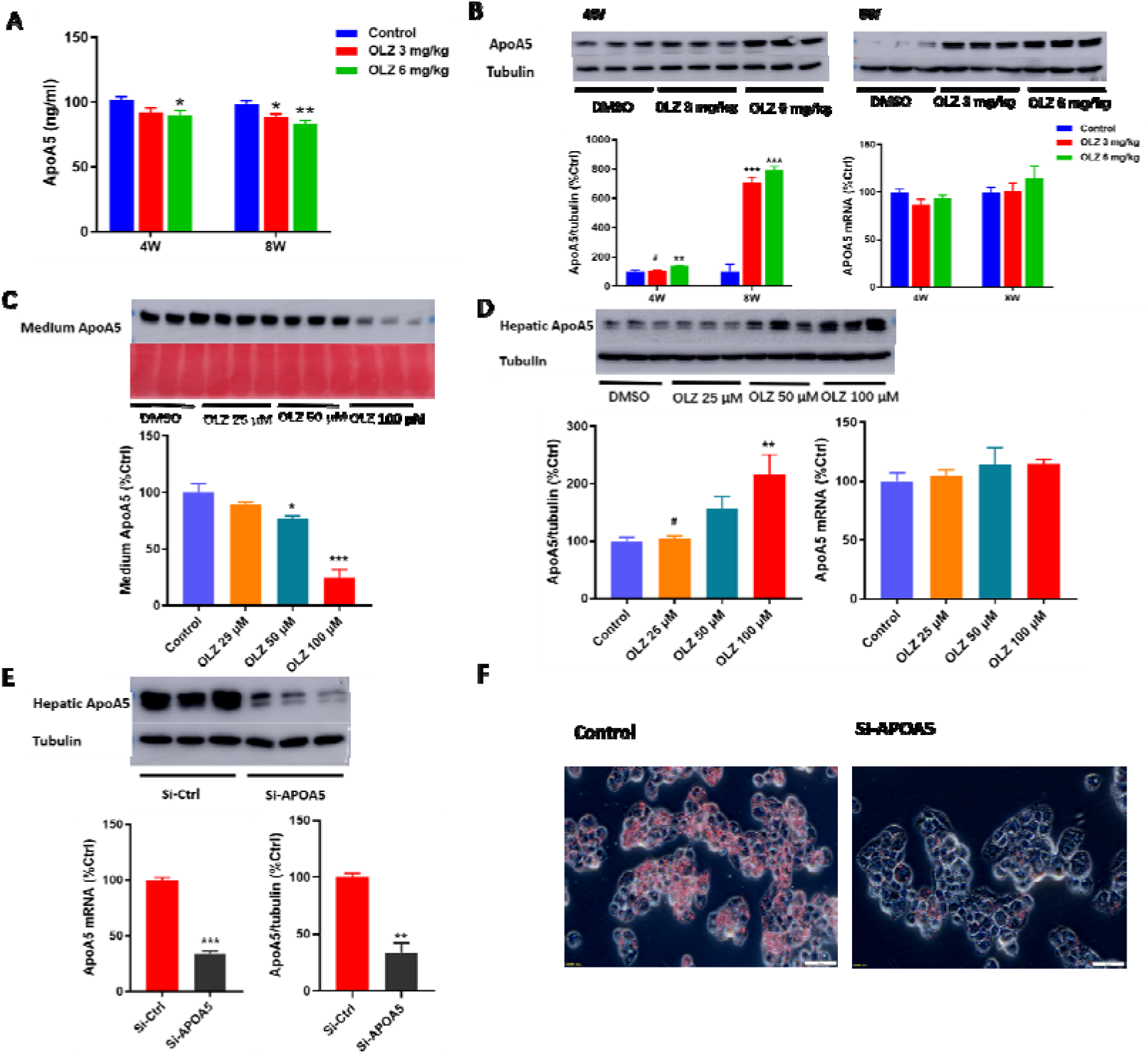
Olanzapine inhibits hepatic apoA5 secretion and exacerbates hepatic steatosis. (**A**) Plasma apoA5 levels in mice, measured by ELISA. (**B**) Hepatic apoA5 expression in the livers of mice based on western blotting and qPCR. (**C**) Western blotting of apoA5 levels in medium:120 μg protein was loaded per lane and loading was confirmed by Ponceau-staining. (**D**) Cellular apoA5 expression in HepG2 cells *in vitro* based on western blotting and qPCR. (**E**) Cellular apoA5 expression in HepG2 cells transfected with *APOA5* siRNA. (**F**) Oil Red O (ORO) staining of HepG2 cells transfected with *APOA5* siRNA. OLZ, olanzapine. Results are shown as the mean ± SEM. *\*p* < 0.05, *\*\*p* < 0.01, *\*\*\*p* < 0.001, versus the control group. #*p* < 0.05 versus the 6 mg/kg group or 100 μM group. Student’s two-tailed t-test and one-way ANOVA plus Tukey’s test were performed for data analysis.

To further verify the involvement of apoA5 in olanzapine-induced NAFLD, we suppressed its expression via the transfection of *APOA5* siRNA into HepG2 cells pre-treated with 100 μM olanzapine. Our ORO staining results indicated that *APOA5* knockdown effectively reversed olanzapine-induced lipid accumulation (Fig 3E). Considering apoA5 as a liver-specific factor secreted into circulation, these *in vivo* and *in vitro* data indicate that the olanzapine-induced reduction in circulating apoA5 should result from impaired secretion instead of the impaired production of hepatic apoA5. As a result, the olanzapine-mediated inhibition of apoA5 secretion would facilitate hepatic apoA5 retention and then result in liver triglyceride accumulation, resulting in hepatic steatosis.

### 3.4. Sortilin is responsible for inhibition of hepatic apoA5 secretion and olanzapine-mediated hepatic steatosis

Given the established role of sortilin in lipoprotein transport/secretion [28] and NAFLD pathogenesis [30–32], as well as the confirmed binding of sortilin to apoA5 *in vitro* [33], it is possible that sortilin is implicated in the olanzapine-mediated inhibition of apoA5 secretion. In this study, we found that olanzapine dose-dependently increased the mRNA and protein levels of hepatic sortilin in mice (Fig 4A). Likewise, olanzapine significantly upregulated the mRNA and protein expression of hepatocyte sortilin *in vitro* (Fig 4B).

**Figure 4.**
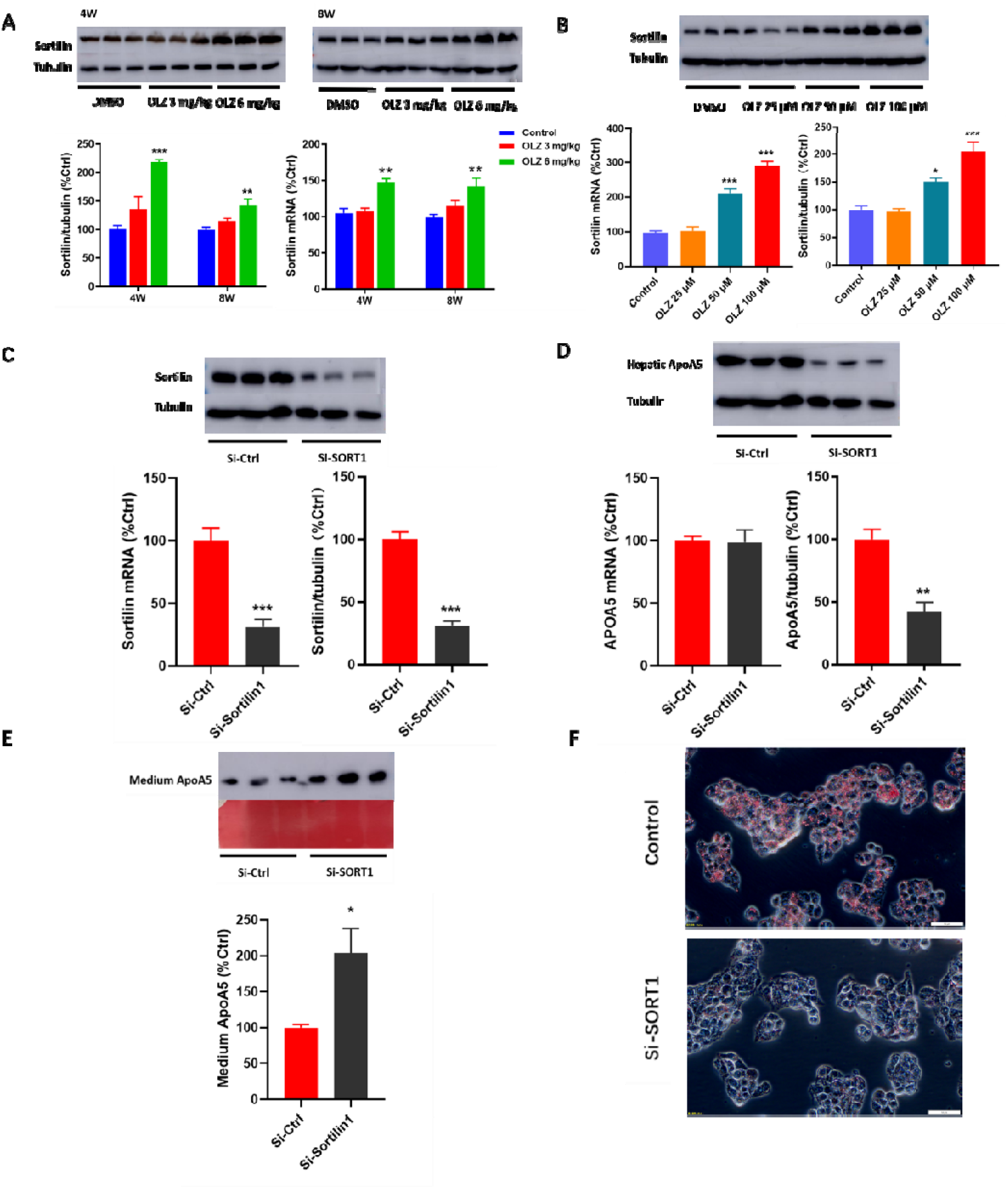
Sortilin participates in the inhibition of apoA5 secretion and olanzapine-mediated hepatic steatosis. (**A**) Hepatic sortilin expression in mice based on western blotting and qPCR. (**B**) Sortilin expression in HepG2 cells based on western blotting and qPCR. (**C**) Sortilin expression in HepG2 cells transfected with *SORT1* siRNA. (**D**) apoA5 expression in HepG2 cells transfected with *SORT1* siRNA. (**E**) Western blotting of apoA5 levels in medium:120 μg protein was loaded per lane and loading was confirmed by Ponceau-staining. (**F**) Oil Red O (ORO) staining of HepG2 cells transfected with *SORT1* siRNA. OLZ, olanzapine. Results are shown as the mean ± SEM. \**p* < 0.05, \*\**p* < 0.01, \*\*\**p* < 0.001 versus the control group. Student’s two-tailed t-tests and one-way ANOVA plus Tukey’s test were performed for data analysis.

Further, we knocked down *SORT1* by transfecting specific siRNA into HepG2 cells pre-treated with 100 μM olanzapine (Fig 4C). Interestingly, *SORT1* knockdown remarkably reduced apoA5 protein levels in cells without affecting *APOA5* mRNA (Fig 4D). In contrast, *SORT1* siRNA increased apoA5 protein levels in the medium (Fig 4E). In addition, ORO staining analysis showed that *SORT1* knockdown effectively attenuated hepatocyte steatosis, characterized by substantially reduced intracellular lipid droplets in these liver cells *in vitro* (Fig 4F). Thus, our findings show that *SORT1* knockdown facilitates the secretion of hepatocyte apoA5, leading to the marked alleviation of hepatic steatosis.

## 4. Discussion

Olanzapine is widely used in clinical practice for patients with schizophrenia because it has fewer extrapyramidal side effects and better therapeutic efficiency [39]. However, the long-term use of olanzapine leads to metabolic syndrome (including obesity, diabetes, and hyperlipidemia), which is associated with an increased risk of NAFLD [39]. The role of olanzapine in the pathogenesis of NAFLD has been established in several animal studies [12,18]. Likewise, our study also confirmed that olanzapine treatment leads to NAFLD development, consistent with previous findings [12,18].

To investigate the mechanism underlying olanzapine-induced NAFLD, apoA5 was considered a candidate factor in this study, which is based on the fact that it is a key regulator of triglyceride metabolism. As stated, disturbances in triglyceride metabolism are the cornerstone of NAFLD pathogenesis. A substantial body of evidence suggests that apoA5 is a negative regulator of circulating triglyceride levels [11, 41, 42]. Several genetic studies have identified *APOA5* gene polymorphisms as a contributing factor to olanzapine-associated dyslipidemia [20, 21]. Additionally, we previously demonstrated that metformin, a drug used for olanzapine-induced metabolic syndrome [43], ameliorates obesity-associated hypertriglyceridemia in mice via the apoA5 pathway [44]. Similarly, this study also found that olanzapine treatment decreased plasma apoA5 levels in mice, with an elevation in plasma triglyceride levels, suggesting that apoA5 is implicated in olanzapine-related hypertriglyceridemia.

However, apoA5 also has a well-documented role in the pathogenesis of NAFLD. Interestingly, its role seems paradoxical in extra- and intrahepatic triglyceride metabolism; that is, apoA5 reduces plasma triglyceride levels and increases hepatic triglyceride contents [42]. In the liver, apoA5 facilitates the biosynthesis of hepatic lipid droplets that promote the development of NAFLD [23, 24, 27]. In this study, olanzapine remarkably increased hepatic apoA5 protein levels and concomitantly resulted in deteriorated liver function and exacerbated hepatic steatosis in mice. Consistently, olanzapine markedly enhanced apoA5 protein levels in hepatocytes *in vitro*, resulting in hepatocyte steatosis, characterized by a substantial elevation of intracellular lipid droplets. In contrast, *APOA5* knockdown effectively ameliorated olanzapine-mediated hepatocyte steatosis *in vitro*. Therefore, our data are the first to establish the role of apoA5 in olanzapine-induced NAFLD.

Considering apoA5 as a liver-specific factor [11, 12], it was initially speculated that decreased hepatic production is attributable to reduced circulating apoA5 levels in these mice. Unexpectedly, we found that olanzapine led to an elevation (not reduction) in apoA5 protein levels in mouse livers and *in vitro* hepatocytes, whereas this drug had no effect on hepatic *APOA5* mRNA levels *in vivo* and *in vitro*. This suggests that reduced circulating apoA5 levels in mice most likely result from posttranslational disturbances in hepatic apoA5 protein turnover. Given that the final plasma concentration of apoA5 depends on both the production and secretion of hepatic apoA5, we hypothesize that impaired apoA5 secretion, rather than impeded apoA5 production (because olanzapine had no observed effect on *APOA5* mRNA expression in our *in vivo* and *in vitro* studies), is responsible for olanzapine-induced NAFLD. According to this, olanzapine inhibits hepatic apoA5 secretion resulting in the retention of this protein in hepatocytes, which ultimately leads to the development of NAFLD.

Of note, our data are partly inconsistent with previous findings indicating that olanzapine-induced dyslipidemia is strongly associated with obesity, resulting from changes in diet (high-fat diet) and lifestyle (less exercise) in patients with schizophrenia [45]. Indeed, the body weight gain of mice in our study did not reach statistical significance, possibly due to the short duration of the study (≤ 8 weeks), which is consistent with results of previous data [46]. Nevertheless, we demonstrated that olanzapine treatment successfully led to hepatic steatosis before body weight gains in rodents. Thus, our findings indicate an alternative metabolic mechanism underlying olanzapine-induced NAFLD; that is, olanzapine, independent of obesity due to overnutrition, directly leads to NAFLD by inhibiting hepatic apoA5 secretion.

The role of sortilin in NAFLD has been confirmed by previous studies, where high *SORT1* expression was found to promote NAFLD pathogenesis [30], but *Sort1* knockout attenuates hepatic steatosis in mice [31, 32]. Recent data showed that sortilin attenuates the secretion of hepatic apolipoprotein B [47], another liver-synthesized and liver-secreted protein, occurring primarily at the posttranslational level [48]. Remarkably, sortilin has been reported to bind apoA5, which is involved in the receptor-mediated endocytosis of apoA5 in human embryonic kidney cells *in vitro* [33], suggesting the possible interaction between apoA5 and sortilin in liver cells. Therefore, the contribution of sortilin to the olanzapine-induced inhibition of apoA5 secretion was further investigated. In this study, we observed that olanzapine enhanced sortilin expression *in vivo* and *in vitro*, which was associated with increased hepatic apoA5 levels and then exacerbated hepatic steatosis. Conversely, *SORT1* knockdown abolished the effect of olanzapine on apoA5 upregulation and lipid deposition in hepatocytes *in vitro*. Interestingly, *SORT1* knockdown resulted in an intra-hepatocyte reduction and an extra-hepatocyte (medium) elevation in apoA5 protein levels, without the observed effect of *SORT1* knockdown on intra-hepatocyte *APOA5* mRNA expression, which suggests that sortilin has an inhibitory effect on hepatocyte apoA5 secretion, rather than hepatocyte apoA5 production. Therefore, our study implied a role for sortilin-mediated inhibition of hepatocyte apoA5 secretion in the pathogenesis of olanzapine-induced NAFLD.

Here, we propose a novel mechanism underlying olanzapine-induced NAFLD (**Fig. 5**). Physiologically, apoA5 is synthesized within the endoplasmic reticulum in hepatocytes, transported to the Golgi body, and efficiently secreted into the blood, thereby orchestrating triglyceride homeostasis and preventing hepatic steatosis. However, olanzapine treatment enhances hepatic sortilin expression, leading to the inefficient secretion and subsequent excessive retention of apoA5 in the liver. As a result, intracellular apoA5 drastically increases, and consequently, it stimulates the accumulation of hepatic triglycerides and facilitates the formation of cytosolic lipid droplets. Finally, the liver undergoes steatosis, which results in the development of NAFLD.

**Figure 5.**
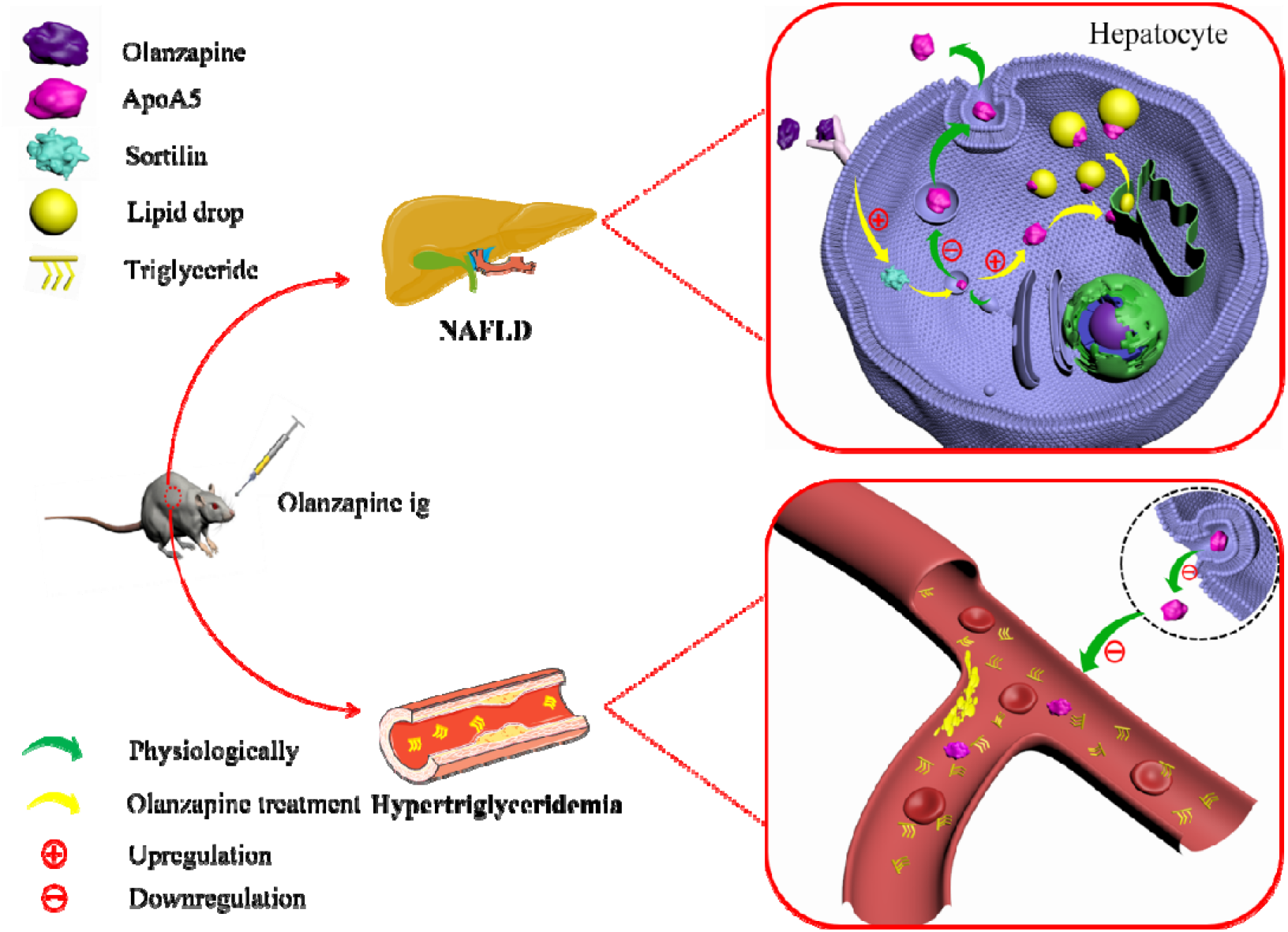
The mechanism underlying olanzapine-induced NAFLD. Physiologically, apoA5 is synthesized and efficiently secreted into the blood, thereby orchestrating extra- and intrahepatic triglyceride metabolism. However, olanzapine treatment pronouncedly enhances hepatic sortilin expression that leads to posttranslational disturbance of hepatic apoA5 protein turnover and inefficient secretion. On the one hand, retention of intracellular apoA5 targeted endoplasmic reticulum and facilitates the formation of cytosolic lipid. Finally, the liver undergo steatosis that confers the development of NAFLD. On the other hand, plasma apoA5 levels in mice drastically decrease with an elevation of plasma triglyceride levels, resulting in olanzapine-related hypertriglyceridemia.

## 5. Conclusion

This study, for the first time, established the role of apoA5 in olanzapine-induced NAFLD. Our findings indicate that olanzapine enhances hepatic sortilin expression, thereby inhibiting hepatic apoA5 secretion, which promotes NAFLD pathogenesis. This suggests that apoA5 might serve as a target for the development of therapeutics for NAFLD in patients with schizophrenia administered olanzapine.

## Abbreviations

ALT: alanine transaminase
ApoA5: apolipoprotein A5
AST: aspartate aminotransferase
DMSO: dimethyl sulfoxide
H&E: hematoxylin and eosin
HDL-C: high-density lipoprotein cholesterol
LDL-C: low-density lipoprotein cholesterol
NAFLD: nonalcoholic fatty liver disease
ORO: Oil Red O

## Contributors

Xiansheng Huang designed the experiments. Wenqiang Zhu and Piaopiao Huang conducted all the experiments and wrote the first draft of the manuscript. Rong Li collected and analysed the data. Fei Luo, Yang Yang, Wen Dai, Wenjing Pei and Li Shen revised the manuscript. All authors contributed to and have approved the final manuscript.

## Data Availability Statement

All data used to support the findings of this study are included within the article.

## Acknowlegement

This study was funded by the National Natural Science Foundation of China (No. 81974281), the Natural Science Foundation of Hunan Province (No.2020JJ2052), National Natural Science Foundation of China (Grant No. 81700999), the Natural Science Foundation of Hunan Province (No.2018JJ3741), XinXin Heart (SIP) Foundation (No.2019-CCA-ACCESS-119). The funding sources had no role in design of the study, analysis of data or writing of the manuscript.

## References

[1] Chalasani N, Younossi Z, Lavine JE, et al. The diagnosis and management of non-alcoholic fatty liver disease: practice Guideline by the American Association for the Study of Liver Diseases, American College of Gastroenterology, and the American Gastroenterological Association. Hepatology. 2012;55(6):2005–2023. doi:10.1002/hep.25762

[2] Bacchi E, Negri C, Targher G, et al. Both resistance training and aerobic training reduce hepatic fat content in type 2 diabetic subjects with nonalcoholic fatty liver disease (the RAED2 Randomized Trial). Hepatology. 2013;58(4):1287–1295. doi:10.1002/hep.26393

[3] Adams LA, Lymp JF, St Sauver J, et al. The natural history of nonalcoholic fatty liver disease: a population-based cohort study. Gastroenterology. 2005;129(1):113–121. doi:10.1053/j.gastro.2005.04.014

[4] European Association for the Study of the Liver (EASL); European Association for the Study of Diabetes (EASD); European Association for the Study of Obesity (EASO). EASL-EASD-EASO Clinical Practice Guidelines for the management of non-alcoholic fatty liver disease. J Hepatol. 2016;64(6):1388–1402. doi:10.1016/j.jhep.2015.11.004

[5] Musil R, Obermeier M, Russ P, et al. Weight gain and antipsychotics: a drug safety review. Expert Opin Drug Saf. 2015;14(1):73–96. doi:10.1517/14740338.2015.974549

[6] Wu RR, Zhao JP, Liu ZN, et al. Effects of typical and atypical antipsychotics on glucose-insulin homeostasis and lipid metabolism in first-episode schizophrenia. Psychopharmacology (Berl). 2006;186(4):572–578. doi:10.1007/s00213-006-0384-5

[7] Wu RR, Zhao JP, Jin H, et al. Lifestyle intervention and metformin for treatment of antipsychotic-induced weight gain: a randomized controlled trial. JAMA. 2008;299(2):185–193. doi:10.1001/jama.2007.56-b

[8] Ballon JS, Pajvani U, Freyberg Z, et al. Molecular pathophysiology of metabolic effects of antipsychotic medications. Trends Endocrinol Metab. 2014;25(11):593–600. doi:10.1016/j.tem.2014.07.004

[9] Liu X, Lian J, Hu CH, et al. Betahistine co-treatment ameliorates dyslipidemia induced by chronic olanzapine treatment in rats through modulation of hepatic AMPK α-SREBP-1 and PPAR α-dependent pathways. Pharmacol Res. 2015;100:36–46. doi:10.1016/j.phrs.2015.07.023

[10] Lin MJ, Dai W, Scott MJ, et al. Metformin improves nonalcoholic fatty liver disease in obese mice via down-regulation of apolipoprotein A5 as part of the AMPK/LXRα signaling pathway. Oncotarget. 2017;8(65):108802–108809. Published 2017 Oct 30. doi:10.18632/oncotarget.22163

[11] Pennacchio LA, Olivier M, Hubacek JA, et al. An apolipoprotein influencing triglycerides in humans and mice revealed by comparative sequencing. Science. 2001;294(5540):169–173. doi:10.1126/science.1064852

[12] van der Vliet HN, Sammels MG, Leegwater AC, et al. Apolipoprotein A-V: a novel apolipoprotein associated with an early phase of liver regeneration. J Biol Chem. 2001;276(48):44512–44520. doi:10.1074/jbc.M106888200

[13] Rensen PC, van Dijk KW, Havekes LM. Apolipoprotein AV: low concentration, high impact. Arterioscler Thromb Vasc Biol. 2005;25(12):2445–2447. doi:10.1161/01.ATV.0000193889.65915.f9

[14] Baroukh N, Bauge E, Akiyama J, et al. Analysis of apolipoprotein A5, c3, and plasma triglyceride concentrations in genetically engineered mice. Arterioscler Thromb Vasc Biol. 2004;24(7):1297–1302. doi:10.1161/01.ATV.0000130463.68272.1d

[15] Lookene A, Beckstead JA, Nilsson S, et al. Apolipoprotein A-V-heparin interactions: implications for plasma lipoprotein metabolism. J Biol Chem. 2005;280(27):25383–25387. doi:10.1074/jbc.M501589200

[16] Mansouri RM, Baugé E, Gervois P, et al. Atheroprotective effect of human apolipoprotein A5 in a mouse model of mixed dyslipidemia. Circ Res. 2008;103(5):450–453. doi:10.1161/CIRCRESAHA.108.179861

[17] Grosskopf I, Shaish A, Afek A, et al. Apolipoprotein A-V modulates multiple atherogenic mechanisms in a mouse model of disturbed clearance of triglyceride-rich lipoproteins. Atherosclerosis. 2012;224(1):75–83. doi:10.1016/j.atherosclerosis.2012.04.011

[18] Brautbar A, Covarrubias D, Belmont J, et al. Variants in the APOA5 gene region and the response to combination therapy with statins and fenofibric acid in a randomized clinical trial of individuals with mixed dyslipidemia. Atherosclerosis. 2011;219(2):737–742. doi:10.1016/j.atherosclerosis.2011.08.015

[19] Huang XS, Zhao SP, Bai L, et al. Atorvastatin and fenofibrate increase apolipoprotein AV and decrease triglycerides by up-regulating peroxisome proliferator-activated receptor-alpha. Br J Pharmacol. 2009;158(3):706–712. doi:10.1111/j.1476-5381.2009.00350.x

[20] Smith RC, Segman RH, Golcer-Dubner T, et al. Allelic variation in ApoC3, ApoA5 and LPL genes and first and second generation antipsychotic effects on serum lipids in patients with schizophrenia. Pharmacogenomics J. 2008;8(3):228–236. doi:10.1038/sj.tpj.6500474

[21] Hong CJ, Chen TT, Bai YM, et al. Impact of apolipoprotein A5 (APOA5) polymorphisms on serum triglyceride levels in schizophrenic patients under long-term atypical antipsychotic treatment. World J Biol Psychiatry. 2012;13(1):22–29. doi:10.3109/15622975.2010.551543

[22] Ress C, Moschen AR, Sausgruber N, et al. The role of apolipoprotein A5 in non-alcoholic fatty liver disease. Gut. 2011;60(7):985–991. doi:10.1136/gut.2010.222224

[23] Feng Q, Baker SS, Liu W, et al. Increased apolipoprotein A5 expression in human and rat non-alcoholic fatty livers. Pathology. 2015;47(4):341–348. doi:10.1097/PAT.0000000000000251

[24] Shu X, Nelbach L, Ryan RO, Forte TM. Apolipoprotein A-V associates with intrahepatic lipid droplets and influences triglyceride accumulation. Biochim Biophys Acta. 2010;1801(5):605–608. doi:10.1016/j.bbalip.2010.02.004

[25] Shu X, Chan J, Ryan RO, Forte TM. Apolipoprotein A-V association with intracellular lipid droplets. J Lipid Res. 2007;48(7):1445–1450. doi:10.1194/jlr.C700002-JLR200

[26] Shu X, Ryan RO, Forte TM. Intracellular lipid droplet targeting by apolipoprotein A-V requires the carboxyl-terminal segment. J Lipid Res. 2008;49(8):1670–1676. doi:10.1194/jlr.M800111-JLR200

[27] Gao X, Forte TM, Ryan RO. Influence of apolipoprotein A-V on hepatocyte lipid droplet formation. Biochem Biophys Res Commun. 2012 Oct 19;427(2):361–5. doi:10.1016/j.bbrc.2012.09.065. Epub 2012 Sep 18. PMID: 23000161; PMCID: PMC3492953.

[28] Strong A, Rader DJ. Sortilin as a regulator of lipoprotein metabolism. Curr Atheroscler Rep. 2012;14(3):211–218. doi:10.1007/s11883-012-0248-x

[29] Strong A, Patel K, Rader DJ. Sortilin and lipoprotein metabolism: making sense out of complexity. Curr Opin Lipidol. 2014;25(5):350–357. doi:10.1097/MOL.0000000000000110

[30] Fisher EA, Ginsberg HN. Complexity in the secretory pathway: the assembly and secretion of apolipoprotein B-containing lipoproteins. J Biol Chem. 2002;277(20):17377–17380. doi:10.1074/jbc.R100068200

[31] Rabinowich L, Fishman S, Hubel E, et al. Sortilin deficiency improves the metabolic phenotype and reduces hepatic steatosis of mice subjected to diet-induced obesity. J Hepatol. 2015;62(1):175–181. doi:10.1016/j.jhep.2014.08.030

[32] Li J, Wang Y, Matye DJ, et al. Sortilin 1 Modulates Hepatic Cholesterol Lipotoxicity in Mice via Functional Interaction with Liver Carboxylesterase 1. J Biol Chem. 2017;292(1):146–160. doi:10.1074/jbc.M116.762005

[33] Nilsson SK, Christensen S, Raarup MK, et al. Endocytosis of apolipoprotein A-V by members of the low density lipoprotein receptor and the VPS10p domain receptor families. J Biol Chem. 2008;283(38):25920–25927. doi:10.1074/jbc.M802721200

[34] Cope MB, Jumbo-Lucioni P, Walton RG, et al. No effect of dietary fat on short-term weight gain in mice treated with atypical antipsychotic drugs. Int J Obes (Lond). 2007;31(6):1014–1022. doi:10.1038/sj.ijo.0803533

[35] Su Y, Liu X, Lian J, et al. Epigenetic histone modulations of PPARγ and related pathways contribute to olanzapine-induced metabolic disorders. Pharmacol Res. 2020;155:104703. doi:10.1016/j.phrs.2020.104703

[36] Albaugh VL, Judson JG, She P, et al. Olanzapine promotes fat accumulation in male rats by decreasing physical activity, repartitioning energy and increasing adipose tissue lipogenesis while impairing lipolysis. Mol Psychiatry. 2011;16(5):569–581. doi:10.1038/mp.2010.33

[37] Yang W, Jiang C, Wang Z, et al. Cyclocarya paliurus extract attenuates hepatic lipid deposition in HepG2 cells by the lipophagy pathway. Pharm Biol. 2020;58(1):838–844. doi:10.1080/13880209.2020.1803365

[38] Chong BF, Murphy JE, Kupper TS, et al. E-selectin, thymus- and activation-regulated chemokine/CCL17, and intercellular adhesion molecule-1 are constitutively coexpressed in dermal microvessels: a foundation for a cutaneous immunosurveillance system. J Immunol. 2004;172(3):1575–1581. doi:10.4049/jimmunol.172.3.1575

[39] Leucht S, Cipriani A, Spineli L, et al. Comparative efficacy and tolerability of 15 antipsychotic drugs in schizophrenia: a multiple-treatments meta-analysis [published correction appears in Lancet. 2013 Sep 14;382(9896):940]. Lancet. 2013;382(9896):951–962. doi:10.1016/S0140-6736(13)60733-3

[40] Gluchowski NL, Becuwe M, Walther TC, et al. Lipid droplets and liver disease: from basic biology to clinical implications. Nat Rev Gastroenterol Hepatol. 2017;14(6):343–355. doi:10.1038/nrgastro.2017.32

[41] Zhao SP, Hu S, Li J, et al. Association of human serum apolipoprotein A5 with lipid profiles affected by gender. Clin Chim Acta. 2007;376(1-2):68–71. doi:10.1016/j.cca.2006.07.014

[42] Forte TM, Ryan RO. Apolipoprotein A5: Extracellular and Intracellular Roles in Triglyceride Metabolism. Curr Drug Targets. 2015;16(12):1274–1280. doi:10.2174/1389450116666150531161138

[43] Wu RR, Zhang FY, Gao KM, et al. Metformin treatment of antipsychotic-induced dyslipidemia: an analysis of two randomized, placebo-controlled trials. Mol Psychiatry. 2016;21(11):1537–1544. doi:10.1038/mp.2015.221

[44] Li R, Chen LZ, Zhao W, et al. Metformin ameliorates obesity-associated hypertriglyceridemia in mice partly through the apolipoprotein A5 pathway. Biochem Biophys Res Commun. 2016;478(3):1173–1178. doi:10.1016/j.bbrc.2016.08.087

[45] Alberti KG, Eckel RH, Grundy SM, et al. Harmonizing the metabolic syndrome: a joint interim statement of the International Diabetes Federation Task Force on Epidemiology and Prevention; National Heart, Lung, and Blood Institute; American Heart Association; World Heart Federation; International Atherosclerosis Society; and International Association for the Study of Obesity. Circulation. 2009;120(16):1640–1645. doi:10.1161/CIRCULATIONAHA.109.192644

[46] Albaugh VL, Henry CR, Bello NT, et al. Hormonal and metabolic effects of olanzapine and clozapine related to body weight in rodents. Obesity (Silver Spring). 2006;14(1):36–51. doi:10.1038/oby.2006.6

[47] Strong A, Ding Q, Edmondson AC, et al. Hepatic sortilin regulates both apolipoprotein B secretion and LDL catabolism. J Clin Invest. 2012;122(8):2807–2816. doi:10.1172/JCI63563

[48] Nielsen MS, Jacobsen C, Olivecrona G, et al. Sortilin/neurotensin receptor-3 binds and mediates degradation of lipoprotein lipase. J Biol Chem. 1999;274(13):8832–8836. doi:10.1074/jbc.274.13.8832

